# Aldh2 is a lineage-specific metabolic gatekeeper in melanocyte stem cells

**DOI:** 10.1101/2021.09.23.461061

**Authors:** Hannah Brunsdon, Alessandro Brombin, Samuel Peterson, John H. Postlethwait, E. Elizabeth Patton

## Abstract

Melanocyte stem cells (McSCs) in zebrafish serve as an on-demand source of melanocytes during growth and regeneration, but metabolic programs associated with their activation and regenerative processes are not well known. Here, using live imaging coupled with scRNA-sequencing, we discovered that quiescent McSCs during regeneration activate a dormant embryonic neural crest transcriptional program followed by an aldehyde dehydrogenase (Aldh) 2 metabolic switch to generate progeny. Unexpectedly, while ALDH2 is well known for its aldehyde clearing mechanisms we find that in regenerating McSCs, Aldh2 activity is required to generate formate – the one-carbon (1C) building block for nucleotide biosynthesis – through formaldehyde metabolism. Consequently, we find that disrupting the 1C cycle with low-doses of methotrexate caused melanocyte regeneration defects. In the absence of Aldh2, we find that purines (but not pyrimidines) are the metabolic end product sufficient for activated McSCs to generate progeny. Together, our work reveals McSCs undergo a two-step cell state transition during regeneration, and that the reaction products of Aldh2 enzymes have tissue-specific stem cell functions that meet metabolic demands in regeneration.

**SUMMARY STATEMENT:** In melanocyte regeneration, quiescent McSCs respond by re-expressing a neural crest identity, followed by an Aldh2-dependent metabolic switch to generate progeny.

## INTRODUCTION

Melanocytes are pigment producing cells that provide black-brown pigmentation in the hair, skin and eyes in the animal kingdom. Melanocytes can emerge directly from the neural crest during development, while other melanocytes come from melanocyte stem cells (McSCs), that are also neural crest derived, and that replenish melanocyte populations in the adult (Mort et al. 2015). In mammals, distinct McSC populations serve as reservoirs for melanocytes that pigment the growing hair shaft, or for skin pigmentation in response to UV-irradiation or wound healing (Nishimura et al. 2005; Adameyko et al. 2009; Chou et al. 2013). In zebrafish, nerve-associated McSCs are an on-demand regenerative population for zebrafish at all stages, and the cell-of-origin for multiple pigment cell types as the zebrafish grows to become an adult (Budi et al. 2008; Budi et al. 2011; Dooley et al. 2013; Singh et al. 2016; Brombin et al. 2021). How McSCs respond to regenerative signals to generate melanocytes is a central question for adult stem cell biology, but also for melanoma pathogenesis, which is increasingly understood to re-activate and depend upon melanocyte lineage developmental programs in disease progression (White et al. 2011; Kaufman et al. 2016; Travnickova et al. 2019; Varum et al. 2019; Johansson et al. 2020; Marie et al. 2020; Baggiolini et al. 2021).

Zebrafish are uniquely poised to study stem cells due to their genetic tractability and amenability to advanced imaging, enabling the intricacies of stem cell and developmental lineages to be followed at the single cell resolution in living animals (Kelsh et al. 1996; Owen et al. 2020; Travnickova and Patton 2021). During zebrafish embryonic development, melanocytes that originate directly from the neural crest generate lateral stripes along the body (Kelsh and Barsh 2011). McSCs that reside at the dorsal root ganglion (DRG) stem cell niche are also established during embryogenesis, are multi-potent, and give rise to glia and multiple pigment cell types that contribute to the adult pigmentation pattern and serve as a source for melanocytes in regeneration (Budi et al. 2008; Hultman et al. 2009; Budi et al. 2011; Johnson et al. 2011; Kelsh and Barsh 2011; Dooley et al. 2013; Singh et al. 2016; Irion and Nusslein-Volhard 2019; Brombin et al. 2021).

Recently, we identified a developmental *tfap2b+* McSC population that we found to be distinct within neural crest and pigment cell lineages (Brombin et al. 2021). Amongst members of the zebrafish Aldehyde dehydrogenase (Aldh) 1 and 2 enzyme family, which are well conserved with analogous human enzymes, we found *aldh2* gene paralogs were expressed in these *tfap2b+* McSCs (**Fig. 1a, S1**). Aldehyde-processing enzymes are viewed as essential clearing agents that rapidly deactivate harmful aldehydes, and also as markers of somatic and cancer stem cell populations (O’Brien et al. 2005; Marcato et al. 2011; Pontel et al. 2015; Garaycoechea et al. 2018). In the bone marrow, two specific enzymes, aldehyde dehydrogenase (ALDH) 2 and alcohol dehydrogenase (ADH) 5, protect hematopoietic stem cells (HSCs) from endogenous formaldehyde accumulation and toxicity (Dingler et al. 2020; Nakamura et al. 2020; Oka et al. 2020; Jung and Smogorzewska 2021; Mu et al. 2021). The importance of aldehyde detoxification in human biology is exemplified by the genetic variants of *ALDH2* in the human population, such as the single nucleotide polymorphism r671 in *ALDH2* (c.1510G>A; p.E504K; ALDH2*2), which confers loss-of-function in 560 million people, mainly of East Asian origin (Chen et al. 2014; Chen et al. 2020). Carriers of the r671 *ALDH2* polymorphism can experience adverse reactions to acetaldehyde from exogenous alcohol consumption and are at risk for a range of diseases including osteoporosis, cardiovascular disease, neurodegeneration, and Fanconi Anemia (Harada et al. 1981; Brooks et al. 2009; Hiura et al. 2010; Takeuchi et al. 2012; Matsuo et al. 2013; Masaoka et al. 2016; Chang et al. 2017).

**Figure 1:**
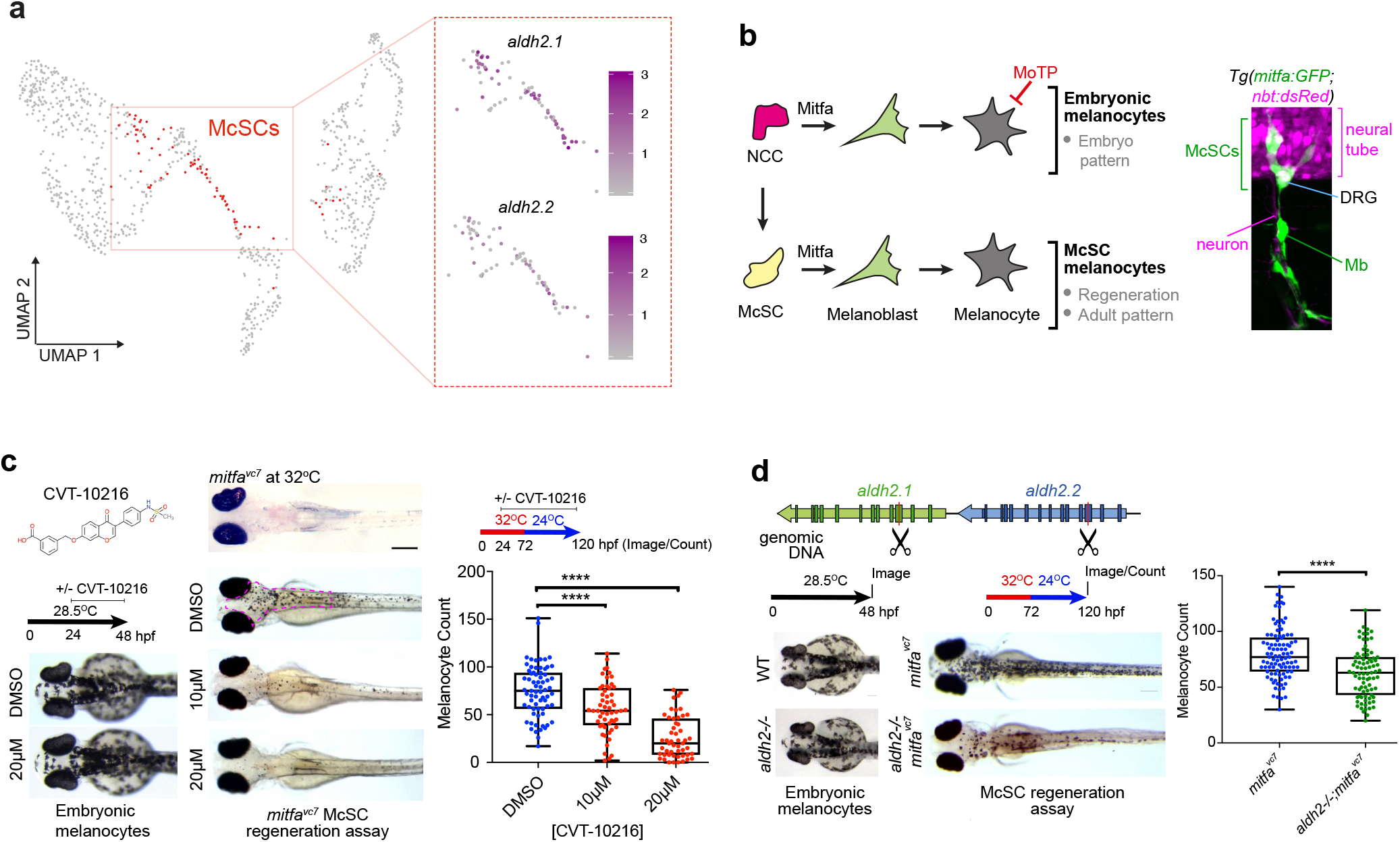
Lineage-specific requirement for Aldh2 in melanocyte regeneration. **a.** UMAP of scRNA-seq data from Brombin et al (2021) with McSCs in red. UMAPs of these isolated McSCs showing log_2_ expression of *aldh2* paralogs with color change from grey (negative) to purple. **b.** Schematic of the melanocyte lineages in zebrafish development with confocal Z-stack depicting McSCs expressing *mitfa:GFP* located at the dorsal root ganglia (DRG) and melanoblasts (Mb) on the motor neurons. Neural tube and DRG are marked by *nbt:dsRed* expression. **c.** Lineage specific Aldh2 activity. Representative images of wild type embryos treated −/− CVT-10216 during development (embryonic melanocytes) or in an McSC regeneration assay. One data point per embryo. Scale bar = 500 μm. **** p<0.0001. One-way ANOVA with Tukey’s multiple comparisons test. N=4. **d.** Schematic of CRISPR-Cas9 strategy to target *aldh2.1* and *aldh2.2* with excision site between Cas9 cut sites (scissor symbols; **see Fig. S1)**. Wild type or *aldh2−/−* embryos in normal development, or a McSC regeneration assay shown. **** p<0.0001. Unpaired two-tailed t-test performed to calculate statistical significance. N=3.

Much of the toxicity from aldehydes can be attributed to metabolites such as acetaldehyde and formaldehyde that cause mutations and chromosomal rearrangements by direct damage to DNA (Pontel et al. 2015; Garaycoechea et al. 2018). Recent work shows that a two-tier protection mechanism in cells defends against aldehyde-induced DNA crosslinks: first, aldehydes are cleared by enzymes, such as ALDH2 and ADH5, and second, replication-coupled DNA damage response pathways repair crosslinks and remove adducts (Langevin et al. 2011; Rosado et al. 2011; Garaycoechea et al. 2012; Pontel et al. 2015; Garaycoechea et al. 2018; Hodskinson et al. 2020). These studies emphasize the nature of aldehyde toxicity and homeostatic clearance, primarily investigated in the hematopoietic stem cell (HSC) compartment. However, other work proposes more varied roles for aldehydes, namely that by-products generated by aldehyde-detoxification enzyme reactions also sustain essential downstream cellular metabolic processes (Jacobson and Bernofsky 1974; Bae et al. 2017; Burgos-Barragan et al. 2017). What is yet unknown is how the reaction products of aldehyde metabolism by ALDH2 contributes to the physiology of specific cells and tissues in processes other than toxicity. Here, we discover a new requirement for Aldh2 dependent metabolism in activated McSCs during regeneration.

## RESULTS

### A lineage-specific function for Aldh2 in melanocyte regeneration

To learn how ALDH2 functions in stem cells other than HSCs and in an intact animal, we set out to study the zebrafish McSC population in melanocyte regeneration. We employed the ALDH2 inhibitor (ALDH2i) CVT-10216 in a melanocyte regeneration assay that is dependent on a temperature sensitive allele (*mitfa^vc7^*) of the master melanocyte transcription factor MITF (Johnson et al. 2011; Zeng et al. 2015). In this model, fish embryos are grown at higher temperatures to deplete Mitfa activity, which prevents embryonic melanocyte development from the neural crest. When the water temperature is lowered to a level permissive for restoring Mitfa activity, melanocytes are regenerated from McSCs (Johnson et al. 2011) **(Fig. 1b)**. In zebrafish embryos grown in the presence of CVT-10216, we did not detect any discernible effects on embryonic melanocyte development. However, melanocyte regeneration from McSCs was significantly delayed in ALDH2i-treated embryos, indicating that Aldh2 has a lineage-specific function in McSCs **(Fig. 1c).**

CVT-10216 is reported to have a >40-fold selectivity for ALDH2 over other ALDH enzymes (Chen et al. 2014), however, to confirm this specificity in zebrafish, we generated an *aldh2.1 - aldh2.2* double mutant line by CRISPR-Cas9, henceforth referred to as *aldh2−/−*. The genetic similarity between these two paralogs made generating specific *aldh2* mutants difficult, so we created a double null mutant instead by designing guide RNAs to excise a large intergenic region between the tandem duplicate genes **(Fig. S1)**. In keeping with our ALDH2i experiments, *aldh2−/−* mutants generated embryonic melanocytes, yet were defective in melanocyte regeneration from the McSC compartment **(Fig. 1d)**. We noticed that after multiple rounds of breeding of our *aldh2−/−* mutants, the melanocyte regeneration phenotype was lessened, coupled with transcriptional upregulation of other *aldh* enzyme family members, suggesting some plasticity in *aldh* expression in regeneration **(Fig. S1)**. To address this, we confirmed the *aldh2−/−* genetic mutant results in *aldh2.1 and aldh2.2* knockdown experiments with morpholino oligonucleotides and once again showed that Aldh2 activity is specifically required in the McSC lineage **(Fig. S1).** Finally, we found that the embryonic melanocytes in *aldh2−/−* mutants were defective for the dopaminergic camouflage response, a neuronally regulated innate behavior, reflecting the function for Aldh2 in dopamine metabolism (Yao et al. 2010). This phenotype recapitulates our previous data with Daidzin, another ALDH2i, and provides confidence that the *aldh2−/−* mutants are defective for Aldh2 activity (Zhou et al. 2012) **(Fig. S1)**.

### Live-imaging captures the McSC requirement for Aldh2 to generate progeny

To investigate whether Aldh2 activity impacts directly upon the McSCs, we employed a *Tg(mitfa:GFP)* transgenic line that was previously shown to mark McSCs (Dooley et al. 2013; Brombin et al. 2021). Following ALDH2i treatment in regenerating embryos, we observed a significant loss of GFP+ expression in McSCs at the niche **(Fig. 2a)**. One interpretation of this result is that McSCs are depleted in the absence of Aldh2. Alternatively, McSCs may be present but expressing only low (or no) *mitfa:GFP* under conditions of ALDH2 inhibition.

**Figure 2:**
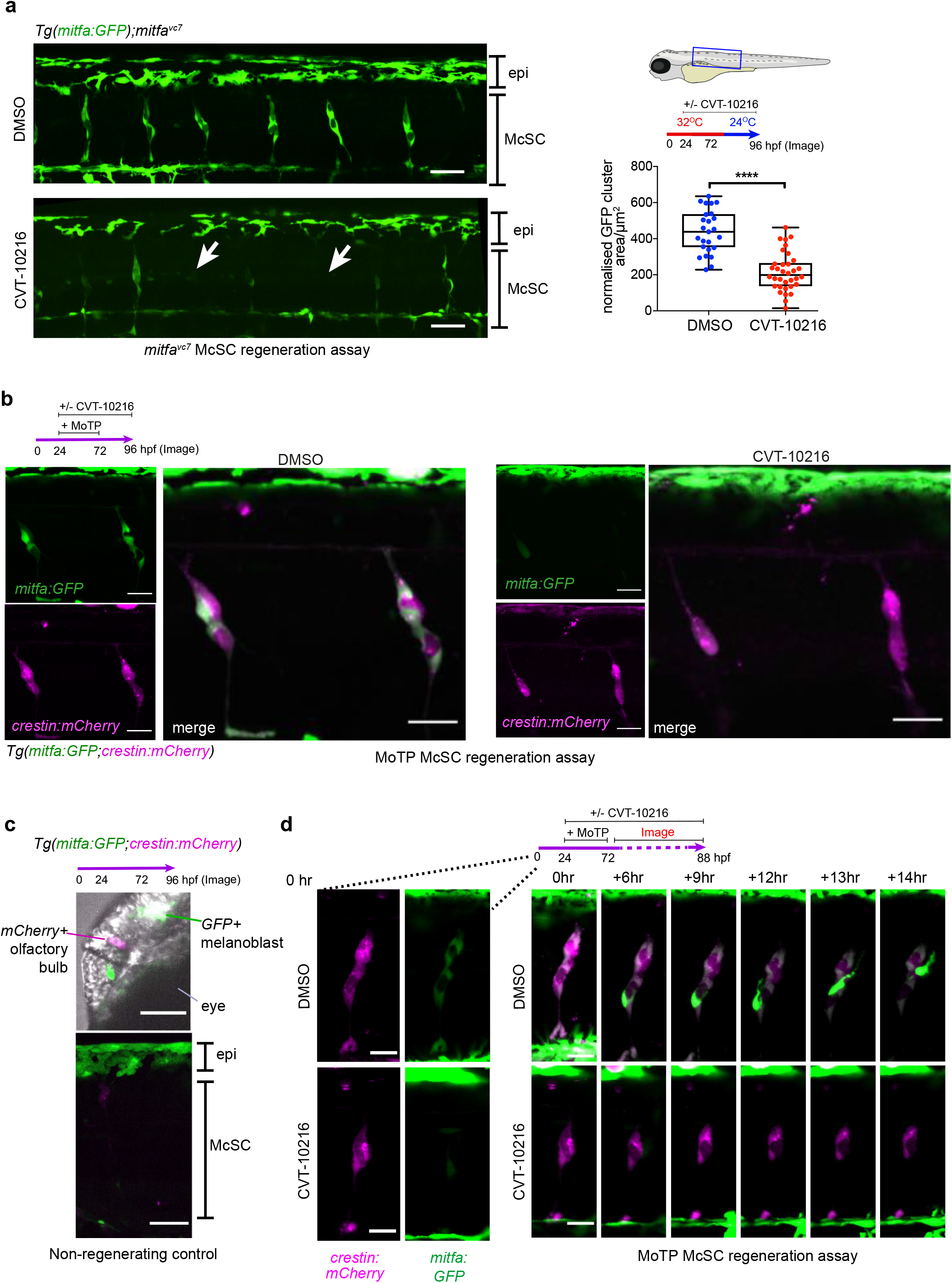
Live-imaging captures the McSC requirement for Aldh2 to generate progeny. **a.** ALDH2 inhibitor (CVT-10216) causes loss of *mitfa:GFP* expression in McSC, while dorsal stripe epithelial (epi) *GFP+* melanoblasts remain. Representative confocal stack images of McSCs at the niche after 24 hours regeneration −/− CVT-10216 treatment. The average *mitfa:GFP* niche area μm^2^/somite was quantified per embryo (one data point). Scale bars are 50 μm, N=3 with >5 embryos imaged per repeat.**** p<0.0001. Unpaired, two-tailed t-test. **b.** McSCs maintain neural crest identity when treated with ALDH2 inhibitor (CVT-10216). Confocal stack images of McSC niches in CVT-10216 treated *Tg(mitfa:GFP;crestin:mCherry)* embryos after 6 hours washout of MoTP. McSCs with very low to no GFP signal are arrowed. N=2, >5 embryos used per condition, representative images shown. Scale bars = 50 μm **c.** 96hpf non-regenerating *Tg(mitfa:GFP;crestin:mCherry)* embryos (same age as **b**) still express *crestin:mCherry* in the olfactory bulb and *mitfa:GFP* in embryonic epithelial melanoblasts (labelled in head, top image, and trunk bottom image), but no longer express these transgenes in McSC niches. Representative images of 3 embryos shown. Scale bars = 50 μm **d.** Time lapse stills of individual regenerating McSCs at the niches. *Tg(mitfa:GFP; crestin:mCherry)* embryos with or without CVT-1016 were imaged from 2 hours post-MoTP washout. In a control embryo, an McSC undergoes cell division and a new *mitfa:GFP*-*high* cell migrates upwards towards the epidermis (see **Movie S1**). In a CVT-10216 treated embryo, *mitfa:GFP* expression is absent, and migration not observed (see **Movie S2**). Scale bars = 20 μm.

In the earliest stages of embryonic development, McSCs that emerge from the neural crest maintain a neural crest identity at the niche, but lose this identity by day 3 (Dooley et al. 2013; Brombin et al. 2021). Given our results in ALDH2i-treated regenerating embryos, we postulated that regenerative (activated) McSCs would re-express neural crest identity markers in addition to *mitfa*. To assess this hypothesis, we employed a double transgenic line *Tg(mitfa:GFP; crestin:mCherry)* in which *mCherry* is expressed from the promoter of the neural crest gene *crestin* (Kaufman et al. 2016; Brombin et al. 2021), and applied this to a second, independent regeneration assay. In this assay, the pro-drug MoTP kills differentiated embryonic melanocytes, and melanocytes are regenerated from the McSC compartment **(Fig. 1b)** (Yang and Johnson 2006). Following MoTP washout, McSCs expressed both mCherry and GFP in control animals **(Fig. 2b)**. McSCs were not detectable in non-regenerating embryos (without MoTP) **(Fig. 2c)**. In regenerating embryos, the intensity of GFP was heterogeneous between McSC clusters, but all McSCs expressed mCherry indicating that McSCs re-express a neural crest identity in regeneration. Upon ALDH2i treatment, and as seen in **Fig. 2a**, we again observed a specific and strong reduction of GFP in McSCs. However, this time, mCherry+ McSCs were still clearly visible. Thus, McSCs re-express a neural crest identity during regeneration and require Aldh2 to increase expression of *mitfa* and generate melanoblasts.

Using live confocal imaging of McSCs to capture this process over time, we performed an MoTP regeneration assay and observed cells expressing high levels of *mitfa:GFP+* emerging from McSCs and migrating dorsally in control embryos **(Fig. 2d; Movie S1)**. In contrast, the McSC niches in ALDH2i-treated embryos had little discernible cell movement, with very little *mitfa:GFP* expression **(Fig. 2d; Movie S2)**. Taken together, these data show that there are at least two distinct cell states within the regenerative McSC niche (*mitfa-low* and *mitfa-high*) and that Aldh2 is required for activated McSCs to increase *mitfa* expression and generate migratory progeny.

### Aldh2, but not Adh5, is required for formaldehyde metabolism in McSCs

To elucidate the mechanism by which Aldh2 affects transitions between cell states in McSCs, we sought to identify its substrate. We reasoned that aldehyde substrates in melanocyte regeneration would be toxic if supplied in excess, and that toxicity would increase in *aldh2−/−* mutant embryos. Therefore, we screened known ALDH2 substrates for sensitivity in zebrafish development overall and specifically in the context of melanocyte regeneration **(Fig. 3a; Fig. S2)**. We found that *aldh2−/−* embryos were resistant to acetaldehyde and propionaldehyde, suggesting an unexpected plasticity in response to these aldehydes. *aldh2−/−* mutants were sensitized to 4-HNE, but this was not specific to the McSC lineage. Importantly, *aldh2−/−* embryos were sensitive to formaldehyde, and notably, low doses of exogenous formaldehyde (that had no other apparent effect on the fish) impaired melanocyte regeneration. This response was significantly stronger in *aldh2−/−* mutants compared to controls **(Fig. 3b, Fig. S2)**. These data indicate that formaldehyde, but not other aldehydes, is an important Aldh2 substrate in the McSC compartment.

**Figure 3:**
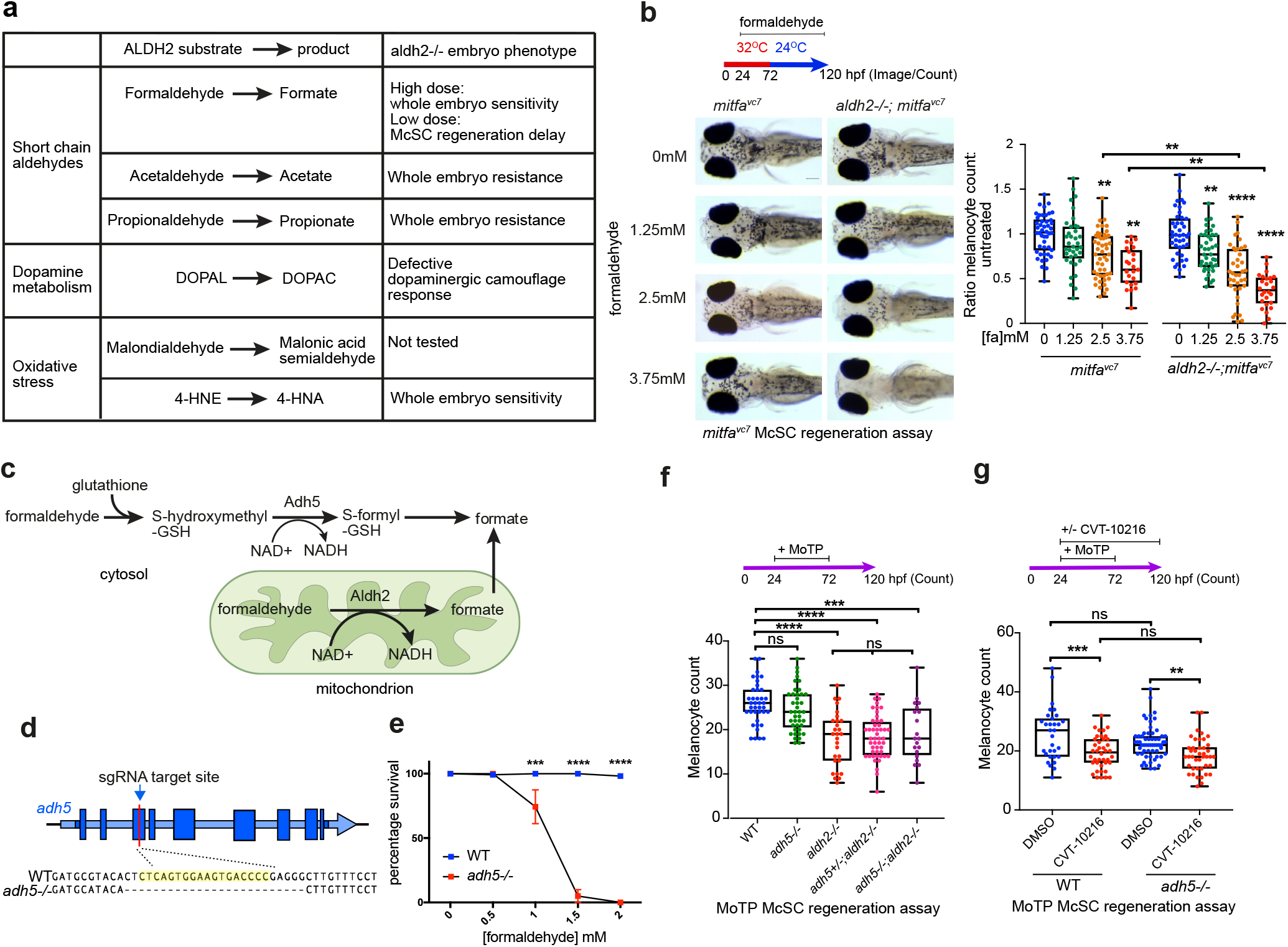
McSCs require Aldh2, but not Adh5, for formaldehyde metabolism. **a.** Table of known ALDH2 substrates and their effects on *aldh2−/−* embryos (See **Fig. S3**). **b.** Melanocyte regeneration is sensitive to formaldehyde and this effect is stronger in *aldh2−/−* mutants. Images and quantification of melanocytes in zebrafish embryos in a *mitfa^vc7^* regeneration assay. Melanocyte counts were normalised to the mean of respective control, each dot represents a single embryo. ** p<0.0021, **** p<0.0001. Ordinary One-way ANOVA with Tukey’s multiple comparisons. N=3. **c.** Schematic diagram of formaldehyde metabolism by Adh5 (cytosol) and Aldh2 (mitochondria). **d.** Schematic diagram showing *adh5−/−* CRISPR-Cas9 mutant, with sgRNA target site in exon 3 and alignment to WT sequence showing a deletion of 25bp. **e.** Sensitivity of *adh5−/−* embryos to increasing concentrations of formaldehyde from 24hpf for 24 hours, and surviving embryos quantified. N=5. ***p<0.0002 **** p<0.0001 Two-way ANOVA with Sidak’s multiple comparisons test, error bars indicate SE. **f.** MoTP regeneration assay on *aldh2−/−*, *adh5−/−* mutant embryos and embryos from an incross of *adh5−/−; aldh2−/−* fish (embryos genotyped after counting). *** p<0.0002, **** p<0.0001, ns not significant. One way ANOVA with Tukey’s multiple comparisons. **g.** MoTP Regeneration assay on wild type and *adh5−/−* mutants treated −/− CVT-10216. N=3. *** p<0.0002, ** p<0.0021, ns: not significant. One way ANOVA with Tukey’s multiple comparisons.

Recent studies show that ALDH2 and ADH5 function together to clear endogenous formaldehyde during HSC differentiation to prevent immune depletion in mouse and induced pluripotent stem cells (iPSCs), as well as in patients with biallelic *ALDH2* and *ADH5* mutations (Dingler et al. 2020; Oka et al. 2020; Shen et al. 2020; Mu et al. 2021) **(Fig. 3c)**. Mice lacking both ALDH2 and ADH5 develop leukemia and have shorter lifespans, and despite active DNA repair, bone marrow-derived progenitors acquire a formaldehyde-associated mutation signature that resembles the human cancer mutation signatures associated with aging (Dingler et al. 2020). To address if Adh5 can function in melanocyte regeneration and compensate for Aldh2, we generated an *adh5−/−* mutant line by CRISPR-Cas9 **(Fig. 3d)**. We found that the *adh5−/−* mutant was highly sensitive to exogenous formaldehyde treatment, indicating that, like in mammals, formaldehyde is an Adh5 substrate in zebrafish **(Fig. 3e)**. However, *adh5* loss had no effect on melanocyte regeneration, and did not enhance the regeneration defects in *aldh2−/−* mutants or ALDH2i-treated embryos **(Fig. 3f, g)**. Thus, despite the shared formaldehyde substrate and conservation across species, Aldh2 has a unique function for formaldehyde metabolism in McSC differentiation, and Adh5 does not compensate for Aldh2 in this cell lineage.

### scRNA-sequencing reveals Aldh2 is a metabolic gatekeeper for McSCs

Thus far, we had visually captured activated McSCs (*crestin+ mitfa-low*) uncoupled from emerging progeny (*crestin+ mitfa-high*), and discovered a novel role for Aldh2 in this process in metabolizing endogenous formaldehyde in these cells. Next, we went on to investigate the transcriptional signatures of these cell populations by scRNA-sequencing (scRNA-seq) to ascertain how they might be affected by Aldh2 deficiency. To this end, we designed a scRNA-seq analysis of a MoTP melanocyte regeneration experiment in which double transgenic *mitfa:GFP; crestin:mCherry* embryos were treated with DMSO or CVT-10216 **(Fig. 4a)**. We identified 24 clusters of transcriptionally distinct cell populations by comparing the top 30 variably expressed genes, generating UMAPs featuring expression of known lineage-defining NC genes, and mapping the cluster identities from two recent zebrafish scRNA publications onto our data (Saunders et al. 2019; Farnsworth et al. 2020) **(Fig. 4b, c; Fig. S3; Tables S1, 2**).

**Figure 4:**
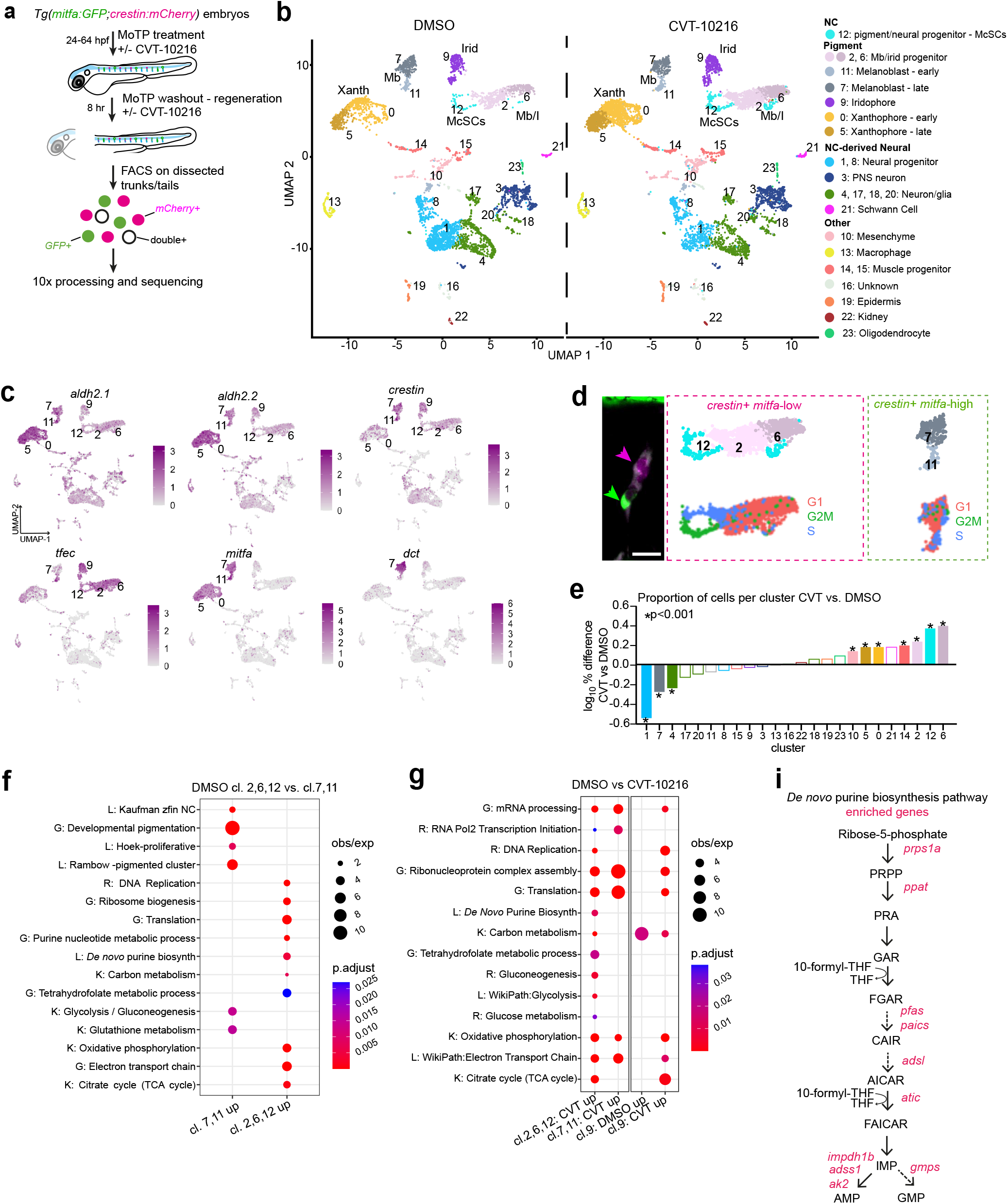
scRNA-seq reveals an Aldh2 metabolic gatekeeper function. **a.** Experimental design for the scRNA-seq experiment to capture the McSCs in regeneration. **b.** UMAPs of *Tg(crestin:mCherry, mitfa:GFP)* positive cells after clustering, split by drug treatment. Mb = melanoblasts, Xanth = xanthophores, Irid = Iridophores. **c.** UMAPs of both DMSO- and CVT-10216 treated cells with color change from grey (negative) to purple based on log_2_ expression of *aldh2.1* and *aldh2.2* in pigment lineages compared to *crestin* (neural crest), *tfec* (melanophore/iridophore progenitors), *mitfa* (early melanoblasts) and *dct* (late melanoblasts). **d.** Proposed relation of imaged McSCs to scRNA-seq clusters, using an example niche from Fig. 1f (scale bar 20 μm). We predict *crestin+ mitfa-high* cells (green arrow/box) are represented in clusters 7, 11, and *crestin+ mitfa-low* cells (magenta arrow/box) are represented in clusters 2,6,12. UMAPs of these clusters (top) and their predicted cell cycle phase (bottom) are shown. **e.** The proportion of total cells within each cluster compared between treatment conditions. The log_10_ percentage difference of numbers of cells in the CVT-10216 treated clusters compared to DMSO equivalents was plotted, with asterisks indicating a significant difference in proportions (Chi squared test). **f.** Dot-plot of pathway analysis showing selection of significantly upregulated GO (G), KEGG (K), Reactome (R) and Literature-based (L) terms in clusters 2,6,12 compared to 7,11, and vice versa. Dot size represents observed/expected ratio, and colour adjusted p-value (Benjamini– Hochberg test). **g.** As **f**, but showing significant enrichment of pathways in CVT-10216 treated cells relative to DMSO from clusters 2,6,12 (*crestin+ mitfa-low)*, clusters 7,11 (*crestin+ mitfa-high)*, and cluster 9 (predicted iridophores). **h.** Schematic diagram of *de novo* purine biosynthesis, with genes encoding enzymes significantly upregulated in the CVT-10216 dataset from **g** and **h** shown in red.

As *crestin:mCherry* is expressed in a wide range of neural crest-derived cell populations (Kaufman et al. 2016), we captured both pigment cell lineages and cells of the neural lineage. The expression of *mitfa* and *dct* in clusters 7 and 11 suggested that these are late and early melanoblast (Mb) populations respectively. Cells in clusters 2, 6 and 12 expressed *crestin*, but low *mitfa*, and contained a mix of markers consistent with McSC identity (Brombin et al. 2021). *aldh2.2* and *aldh2.1* were expressed across multiple clusters but were particularly enriched in regenerating pigment clusters including melanoblasts **(Fig. 4c)**. Relating the above cluster identities to our imaging analyses, we propose that the *crestin+ mitfa-low* McSCs are within clusters 2, 6 and 12 and that the *crestin+ mitfa-high* McSCs and progeny (and any remaining embryonic melanoblasts) are within clusters 7 and 11 **(Fig. 4d)**. The predicted cell cycle phase shows clusters 11 (*mitfa-high*) and 12 (*mitfa-low*) to be in S and G2/M, and may reflect the cycling McSCs we observe during regeneration **(Fig. 2c, 4d)**.

Next, we analyzed the dataset by drug treatment condition. Overall, we found that Aldh2 inhibition did not substantially change cell or cluster identity **(Fig. 4b)**. However, the proportions of cells within some clusters differed significantly between treatment conditions **(Fig. 4e)**. Specifically, we detected a higher proportion of *crestin+ mitfa-low* cells (clusters 2,6,12), and a lower proportion of *crestin+ mitfa*-*high* cells (cluster 7) after ALDH2i. This population shift is consistent with our imaging experiments, in which we detected fewer *mitfa:GFP* expressing cells at the McSC niche **(Fig. 2a-c)**, and suggestive of a block in McSC differentiation.

To understand the physiological and mechanistic implications of the ALDH2-dependent *mitfa-high* to *mitfa-low* McSC transition, we performed differential expression analysis with the control dataset between *crestin+ mitfa-low* cells and *crestin+ mitfa-high* cells **(Table S3)**. *mitfa-high* cells (clusters 7,11) were enriched for pigmentation programs and melanoma-related terms, whereas *mitfa-low* cells (clusters 2,6,12) were enriched for essential metabolic pathways, including the 1C (THF) cycle, the TCA cycle, and *de novo* purine biosynthesis **(Fig. 4f)**, suggesting that regenerative McSCs acquire metabolic requirements distinct from those of melanoblasts.

Next, to understand why McSCs require Aldh2 activity to generate progeny, we performed differential expression analyses between controls and ALDH2i-treated cell populations **(Fig. 4g), Tables S4-6**). Within the ALDH2i treated *crestin+ mitfa-low* cell population, *de novo* purine synthesis was again significantly upregulated **(Fig. 4g, h; Fig S3)**, suggesting that McSCs “blocked” by ALDH2i are starved for purines. We found no ALDH2i-dependent change in *de novo* purine synthesis or glucose metabolism genes in cells from either clusters 7,11 (melanoblast) or another pigment cell cluster requiring purine synthesis for pigmentation (cluster 9; iridophores) (Ng et al. 2009). Therefore, this pattern was specific to *crestin+ mitfa-low* cells and not a general effect of drug treatment. Taken together, these analyses support a mechanism in which regenerative McSCs require Aldh2 for metabolic rewiring to generate progeny.

### Formate, the reaction product of Aldh2-dependent formaldehyde metabolism, promotes McSC transitions

One explanation for the Aldh2-deficient regeneration phenotype is that accumulation of endogenous formaldehyde causes McSC toxicity. However, we believe this to be unlikely given our experimental data; i) our observations while imaging over time showed no evidence of McSC loss, ii) following ALDH2i treatment, *crestin+ mitfa-low* McSCs were present in our scRNA-seq analysis, even at relatively higher numbers, and iii) the McSC block by ALDH2i treatment was reversible following washout **(Fig. S4)**. These findings led us to hypothesize that the reaction products of formaldehyde metabolism are required for timely McSCs differentiation but not for survival **(Fig. 5a)**. To test this hypothesis, we performed a regeneration assay in CVT-10216 treated embryos in the presence or absence of formate, and found that formate supplementation fully restored melanocyte regeneration **(Fig. 5b)**. At the cellular level, formate even fully rescued *crestin+ mitfa-high* expression at the niche site, while having no discernible effect on *crestin+ mitfa-low* cells **(Fig. 5c)**. These results indicate that formate, an Aldh2-dependent reaction product, promotes McSCs to transition from a *mitfa-low* to *mitfa-high* state to generate progeny.

**Figure 5.**
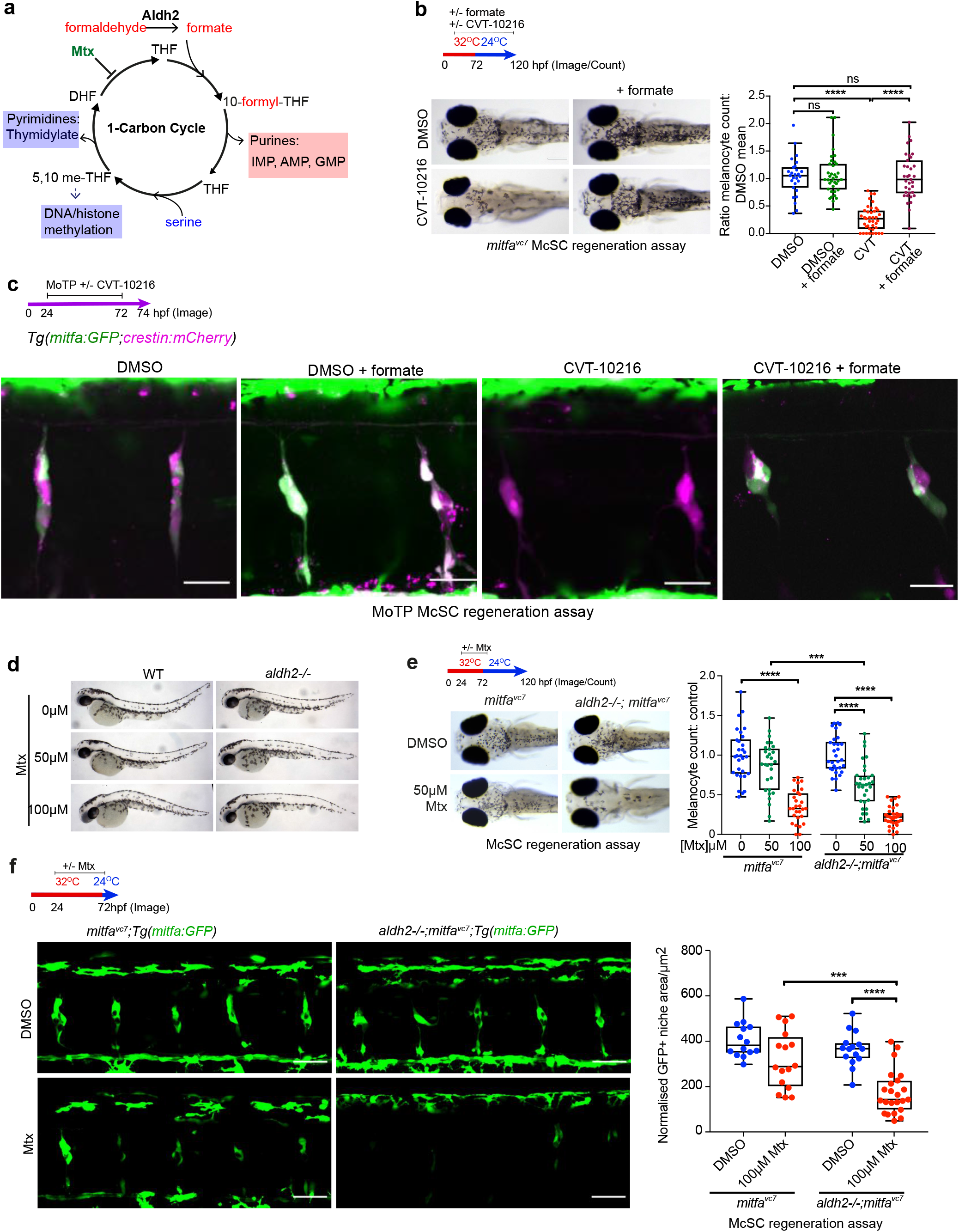
The Aldh2 metabolic reaction product, formate, promotes McSC-derived progeny. **a.** Schematic of 1C metabolism and proposed function for Aldh2 supply of formate through formaldehyde metabolism (based on Burgos-Barragan et al., 2017). Tetrahydrofolate (THF) combines with formate to make 10-formyl-THF, which directly provides two carbons to make purine nucleosides (inosine, adenosine, and guanosine) which can then be converted into nucleotides such as AMP. Serine also donates carbon units to THF as a methyl group to make 5-10,methylene-THF, which in turn donates carbon for DNA/histone methylation, as well in pyrimidine synthesis. **b.** Formate rescues the ALDH2i melanocyte regeneration phenotype. Representative images of a regeneration assay where control or CVT-10216-treated embryos were supplemented with 25 mM sodium formate. P value: **** p<0.0001, ns= not significant. Kruskall-Wallis test with Dunn’s multiple comparisons. N=3. **c.** Formate rescues the McSC Aldh2i differentiation deficiency. An MoTP assay on *Tg(mitfa:GFP;crestin:mCherry)* embryos treated with or without CVT-10216 −/− 25 mM sodium formate from 24hpf. MoTP was washed out at 72hpf, and embryos imaged confocally at 74hpf. N=2, >5 embryos imaged per replicate. Scale bars are 25 μm. Single channel images of *crestin:mCherry* expression (magenta) and *mitfa:GFP* expression (green) are shown alongside merged images. **d.** Mtx treatment has no effect on embryonic melanocytes. Zebrafish embryos (wildtype and *aldh2−/−*) treated with or without Mtx at 24 hpf for 48 hr. N=3. **e.** Melanocyte regeneration is sensitive to disruption of the 1C cycle. Representative images of control and *aldh2−/−* mutants −/− Mtx treatment in a *mitfa^vc7^* regeneration assay are shown. To compare regeneration counts between genotypes, the melanocyte count at each dose was normalised to its respective genotype DMSO control. Each dot represents a single embryo. *** p<0.0002, **** p<0.0001. Ordinary One-way ANOVA performed with Tukey’s multiple comparisons test. N=3. **f.** McSCs are sensitive to disruption of the 1C cycle. Confocal Z-stacks of *mitfa:GFP* McSCs in a *mitfa^vc7^* regeneration assay, in control or *aldh2−/−* embryos treated with or without Mtx. Scale bars are 50 μm. N= 2 biological repeats, with >5 embryos imaged per repeat. Quantification of GFP+ niche area/somite of embryos treated with Mtx is shown. *** p<0.0002, **** p<0.0001. Ordinary One-way ANOVA with Tukey’s multiple comparisons.

### McSCs require a functional 1C cycle

Formate is a carbon donor for the 1C cycle. We found the McSC metabolic switch identified here was reminiscent of cell state transitions reported for naïve to primed murine stem cells that depend on 1C cycling and nucleotide biosynthesis (Chandrasekaran et al. 2017), as well as formate overflow mechanisms that induce a metabolic shift from low to high adenine nucleotide levels in human cancer cell lines and mouse cancer models (Oizel et al. 2020). Indeed, 1C metabolism, compartmentalized within different cell types and organs, is becoming more broadly recognized as a physiological process impacting on cell states and associated with disease (Ducker and Rabinowitz 2017). Taken together, our data suggest that regenerative McSCs depend on 1C cycling to transition from a neural crest to a melanoblast cell state.

To test this hypothesis, we used the dihydrofolate reductase inhibitor methotrexate (Mtx) to inhibit 1C metabolism **(Fig. 5a, d)**. Mtx had no effect on the embryonic melanocyte lineage but its inhibitor function was easy to validate in zebrafish embryos; wild-type embryos treated with Mtx lack pigmentation in xanthophores and iridophores, both of which require functional 1C metabolism for pigment synthesis (Ng et al. 2009) **(Fig. 5d, S4)**. In the McSC lineage, we found that Mtx treatment caused melanocyte regeneration defects that were significantly exacerbated in *aldh2−/−* mutants **(Fig. 5e, f)**. These data indicate that zebrafish McSCs have metabolic requirements that require functional 1C metabolism.

### Aldh2-dependent formaldehyde metabolism meets the demand of McSCs for purines

Given the upregulation of *de novo* purine metabolism genes in McSCs and their dependency on 1C metabolism, we next set out to examine purine nucleotide supplementation in regeneration. In the presence of ALDH2i, we found that exogenously provided purine nucleotides rescued the melanocyte regeneration defect in a dose-dependent manner **(Fig. 6a)**. This effect was not simply a consequence of providing embryos with an additional energy source in the form of ATP, because purine ribonucleosides were also capable of rescuing melanocyte regeneration **(Fig. 6b)**. However, pyrimidine supplementation did not rescue melanocyte regeneration, demonstrating that this effect does not reflect a general requirement for all nucleotides. Next, we explored the specificity of this rescue using confocal imaging, and found that purine, but not pyrimidine, supplementation selectively rescued *mitfa:GFP* expression at the McSC niche after ALDH2i treatment **(Fig. 6c, d)**. Hence, McSCs have a specific requirement for Aldh2 to generate progeny, and the end product of Aldh2 formaldehyde metabolism in McSCs is purine nucleotides **(Fig. 6e)**.

**Figure 6:**
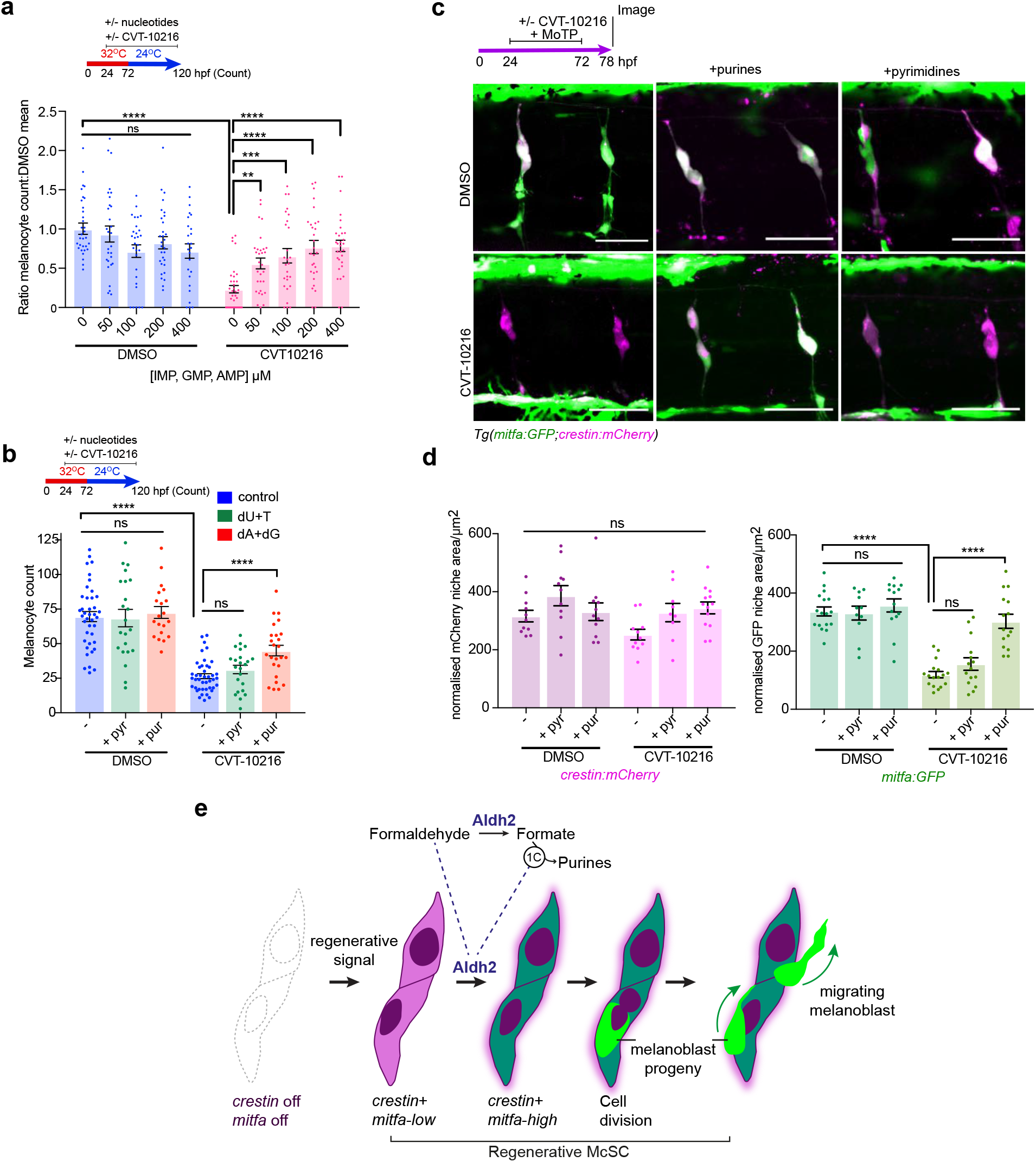
Aldh2 meets the demand of McSCs for purines. **a.** Purine nucleotides rescue Aldh2 deficient melanocyte regeneration. Melanocyte regeneration assay in *mitfa^vc7^* embryos −/− CVT-10216 plus increasing concentrations of purine nucleotides (IMP,GMP and AMP cocktail). N=3. Melanocyte counts are normalized to the untreated control. Each dot represents a single embryo. Error bars represent SE. ** p<0.0021, *** p<0.0002, **** p<0.0001, ns= not significant. Ordinary One-way ANOVA performed with Tukey’s multiple comparisons test with a single pooled variance. **b.** Purine, but not pyrimidine nucleosides, rescue Aldh2 deficient melanocyte regeneration. Melanocyte regeneration assay on *mitfa^vc7^* embryos −/− CVT-10216 and supplemented with deoxyadenosine (dA), deoxguanosine (dG) purine nucleosides, or deoxyuridine (dU) or thymidine (T) pyrimidine nucleosides (200 µM) N=3. **** p<0.0001, ns= not significant. Error bars represent SE. Ordinary One-way ANOVA performed with Tukey’s multiple comparisons test with a single pooled variance. **c.** Purine nucleotides rescue McSC differentiation in ALDH2 inhibitor treated embryos. Representative confocal Z-stacks of *Tg(mitfa:GFP;crestin:mCherry)* embryos treated with MoTP −/− CVT-12016, as well as 400 µM AMP/GMP purine nucleotides, or 400 µM UMP or Thymidine pyrimidine nucleotides. **d.** Quantification of *crestin:mCherry* and *mitfa:GFP* niche areas from **c.** Each dot represents the sum of the GFP or mCherry niche area/ number of somites in view in one embryo. ****:p<0.0001, ns: not significant. Error bars represent SE. Ordinary One-way ANOVA performed with Tukey’s multiple comparisons test with a single pooled variance. **e.** Proposed model for Aldh2 control of the McSC lineage. Regenerating McSCs start expressing *crestin* and low levels of *mitfa*. Next, McSCs increase their metabolic demands for purine nucleotides to express high levels of *mitfa* and generate progeny. This metabolic demand is met by Aldh2, that metabolises endogenous formaldehyde into formate, which is then used in the 1C cycle to fuel the production of purine nucleotides. McSCs undergo cell division to generate progeny, which migrate away from the niche to the epidermis. ALDH2i (CVT-10216) delays the progression of the activated McSC to generate progeny in regeneration.

## DISCUSSION

Understanding McSC responses to regenerative signals is central to the search for druggable targets for regenerative medicine and melanoma therapies **(Patton et al. 2021)**. Here, we coupled single cell RNA-sequencing with live imaging and chemical-genetics in zebrafish McSCs to delineate how quiescent McSCs become activated and then transition to a proliferative state. By screening aldehyde substrates, we find melanocyte regeneration is sensitive to formaldehyde, is independent of *adh5*, and that the reaction product formate is sufficient to rescue Aldh2 deficiency. Thus, we identified an Aldh2-dependent mechanism exerting metabolic control of regeneration in McSCs, distinct from its aldehyde clearing mechanism. 8% of the world’s population carry activity-reducing ALDH2 mutations and the underlying disease mechanism is considered to be elevated cellular toxicity. Thus, identification of an ALDH2-dependent gatekeeper mechanism for a regenerative stem cell response may have important ramifications for carriers of inactivating ALDH2 variants.

We find that regenerative McSCs re-activate a neural crest identity, which is reminiscent of the neural crest and melanocyte developmental states that become reactivated in melanoma disease progression (White et al. 2011; Shakhova et al. 2012; Konieczkowski et al. 2014; Kaufman et al. 2016; Rambow et al. 2018; Travnickova et al. 2019; Varum et al. 2019; Diener and Sommer 2020; Johansson et al. 2020; Marie et al. 2020). Notably, while all cells require nucleotides as fundamental building blocks, and for energy and signaling, the neural crest is especially sensitive to nucleotide depletion, which has direct metabolic consequences in rare disease and melanoma (Sporrij and Zon 2021). For instance, patients with Miller syndrome, a rare genetic neurocristopathy affecting face and limb development, have mutations in dihydroorotate dehydrogenase (DHODH), the rate-limiting enzyme for pyrimidine *de novo* biosynthesis (Ng et al. 2010; Sporrij and Zon 2021). In zebrafish, expression of a neural crest program defines melanoma initiation, and these cancers are sensitive to leflunomide, a DHODH inhibitor (White et al. 2011; Kaufman et al. 2016). Similarly, in mouse, a metabolic gene program driven by the transcription factor Yin Yang 1, a neural crest stem cell regulator, is essential for neural crest lineages - its loss of function causes hypoplasia and prevents initiation of melanoma (Varum et al. 2019). In these contexts, nucleotide sensors may directly influence the transcriptional response, as we and others have shown for the neural crest and McSC differentiation (Johansson et al. 2020; Santoriello et al. 2020).

We were surprised to discover that that regenerative McSCs have a select requirement for purine nucleotides (rather than pyrimidine nucleotides), findings that may point to purine nucleotide functions beyond transcription or DNA replication. For instance, purine nucleotides have an ancient function as neurotransmitters that activate purinergic receptors, and as such can regulate neural stem and progenitor cells, and melanocyte-keratinocyte communication in human skin (Ulrich et al. 2012; Lee et al. 2019). Hence, purine nucleotides could facilitate McSC communication with DRG niche cells (of which we know very little) and with peripheral nerves that are used as migratory routes for melanoblast progenitors (Budi et al. 2011; Dooley et al. 2013). Given that neural crest and McSCs programs re-emerge in melanoma, our findings may be relevant to understanding the metabolic reprogramming in melanomas, such as the dependency on formate metabolism during melanoma metastasis (Piskounova et al. 2015; Fischer et al. 2018).

How stem cells generate progeny is a fundamental question in regenerative medicine. Here, we show that Aldh2-dependent formaldehyde metabolism underlies McSCs metabolic demand for purines to generate progeny. Formaldehyde is abundant in the blood (>40 µM) and can arise from demethylation reactions from histones and nucleic acids (Dingler et al. 2020; Mu et al. 2021). While ALDH2 is often thought of as a protective enzyme, we find no evidence of McSC toxicity in zebrafish with defective Aldh2 activity. Based on our data in **Fig. 3**, we suggest that an unknown endogenous formaldehyde source is active in melanocyte regeneration. Conceptually, our work identifies an unanticipated lineage-specific requirement for Aldh2 in the supply of essential metabolites in McSCs. This could mean that in individuals with inactivating mutations in ALDH2, both aldehyde cytotoxicity and depletion of aldehyde derived metabolites could result in the clinical disease features.

## MATERIALS AND METHODS

### Data and code availability

scRNA-seq experiment data have been submitted to GEO (GSE183868). A private access token is available for reviewers. Previously published sequencing data that was reanalyzed here are available from GEO: GSE131136 (Saunders et al. 2019), NCBI SRA: PRNJNA56410 (Farnsworth et al. 2020), and GEO: GSE178364 (Brombin et al. 2021).

### Fish husbandry, fish lines

Zebrafish were maintained in accordance with UK Home Office regulations, UK Animals (Scientific Procedures) Act 1986, amended in 2013, and European Directive 2010/63/EU under project license 70/8000 and P8F7F7E52. All experiments were approved by the Home Office and AWERB (University of Edinburgh Ethics Committee). Fish stocks used were: wild-type AB, *mitfa^vc7^* (Johnson et al. 2011; Zeng et al. 2015), *Tg(mitfa:GFP)* (Dooley et al. 2013), *Tg(crestin:mCherry)*(Kaufman et al. 2016), *aldh2−/−* (this study), and *adh5−/−* (this study). Combined transgenic and mutant lines were generated by crossing. Adult fish were maintained at ∼28.5°C under 14:10 light-dark cycles. Embryos were kept at either 24°C, 28.5°C or 32°C and staged according to the reference table provided by Kimmel and colleagues (Kimmel et al. 1995).

### Genotyping

Whole embryos or fin clips from adult fish were genotyped by resuspending tissue in DirectPCR® DNA-Tail solution (Viagen), and heating samples to 56°C for 2 hours, then 84°C for 20 minutes. Primers used for genotyping can be found in **Table S7**.

### CRISPR-Cas9 mutant line generation

sgRNAs (**Table S7**) were synthesized using the EnGen® sgRNA Synthesis Kit, *S. pyogenes* (New England Biolabs) according to manufacturer’s instructions. CRISPR-Cas9 knock-out lines were generated as previously described (Sorlien et al. 2018). Briefly, 200 ng/μl sgRNAs targeting exon 3 of *aldh2.1* (GCCAGAGATGCCTTTAAGCT) and exon 3 of *aldh2.2* (GCCAGAGATGCCTTTAAGCT) were co-injected with Cas9 mRNA into zebrafish embryos at the 1 cell stage. An allele was recovered which was the result of a large deletion between *aldh2.1* and *aldh2.2*, creating a gene fusion and single base-pair insertion at the fusion site. This introduced an adjacent frameshift mutation and premature stop codon. 200 ng/μl sgRNA targeting exon 3 of *adh5* (CTCAGTGGAAGTGACCCCGAG) was co-injected with recombinant 300 ng/μl Cas9 protein (SBI). These F0 fish were raised to adulthood, and outcrossed with WT fish to obtain progeny that were screened for presence of indels through PCR amplification of a 600bp region surrounding the target site, and digestion of the amplicon using T7 endonuclease (New England Biolabs). Outcrossed F1 fish that contained a 25bp deletion were isolated and raised to adulthood.

### Morpholino injection

Standard control morpholinos and translation blocking morpholinos were sourced from Genetools LLC, based off previously published sequences for *aldh2.1* (ZDB-MRPHLNO-120517-2) and *aldh2.2* (ZDB-MRPHLNO-120517-3)(Ma et al. 2010). 2-6 ng of each morpholino was injected into sibling *mitfa^vc7^* embryos at the 1-2 cell stage.

### Imaging

Images of embryos immobilized with MS:222 and 1.5% LMP agarose were acquired using a 20X/0.75 lens on the multimodal Imaging Platform Dragonfly (Andor technologies, Belfast UK) equipped with 405, 445, 488, 514, 561, 640 and 680nm lasers built on a Nikon Eclipse Ti-E inverted microscope body with Perfect focus system (Nikon Instruments, Japan). Data were collected in Spinning Disk 40 μm pinhole mode on the Zyla 4.2 sCMOS camera using a Bin of 1 and no frame averaging using Andor Fusion acquisition software. Z stacks were collected using the Nikon TiE focus drive. Multiple positions were collected using a Sigma-Koki Stage (Nikon Instruments Japan). Data were visualized and analyzed using Imaris (Oxford Instruments, v. 9.7.0) or Image J Fiji software (v. 1.53c).

Whole zebrafish embryos fixed in 4% PFA/PBST were imaged with a Leica MZFLIII fluorescence stereo microscope with a 1x objective fitted with a Qimaging Retiga Exi CCD camera (Qimaging, Surrey, BC, Canada). Image capture was performed using Micromanager (Version 1.4).

To quantify the area of *GFP* or *mCherry*-expressing cells within niches, homozygous *Tg(mitfa:GFP)* fish were outcrossed with non-fluorescent fish to obtain embryos with similar levels of transgene expression. The McSC compartment was imaged at the same magnification, within the same anatomical area, and with consistent laser power and other imaging settings between individual samples and biological replicates. In Fiji, a maximum projection Z-stack of images was cropped to only include McSC compartment cells (typically containing 6-7 compartments per image) and converted to a binary image. Consistent threshold settings were applied, and the total GFP+ area measured in pixels^2^ and divided by the number of somites visible in the field of view.

### Melanocyte regeneration assays

If using the *mitfa^vc7^* regeneration model line, embryos were kept in a 32°C incubator from 0-72hpf to repress the developmental melanocyte lineage. Embryos were then moved to a 24°C incubator to allow regeneration over a period of 48 hours. When using chemical methods for regeneration, 150 μM 4-(4-Morpholinobutylthio)phenol (MoTP) (Sigma) was added to embryos kept at 28.5°C from 24hpf onwards. MoTP was washed out to allow regeneration between 72 and 120 hpf. After fixation, embryos were imaged and melanocytes counted within a set region with the Image J CellCounter plugin.

### Camouflage response assays

The camouflage response assay was performed as described previously (Zhou et al. 2012). 5 dpf wild type or *aldh2−/−* mutant embryos were placed in a dark place for 15 minutes to standardize their light exposure. These embryos were split into cohorts which were either placed under a lamp or kept in the dark for 1.5 hours. The embryos were then moved to the opposite light condition for a further 45 minutes, during which time melanin dispersed or contracted depending on light exposure. This was repeated once or twice more when assessing the embryos ability to learn to adapt to changing light conditions. Afterwards, embryos were then briefly anaesthetized in MS-222 and fixed in 4% PFA. Embryos were imaged dorsally at a fixed magnification. Melanin coverage was measured with Image J Fiji, by outlining a predetermined region of the head, converting the image to an 8-bit binary image with a uniform threshold, and then measuring the area of black pixels.

### Small molecule inhibitor and rescue experiments

Unless otherwise stated, 10 μM CVT-10216 (Sigma-Aldrich) or equimolar Dimethyl Sulphoxide DMSO (Sigma-Aldrich) was added to embryos at 24hpf after manual or pronase-assisted (Sigma) dechorionation and refreshed every 24 hours. Embryos were arrayed in 6-well tissue culture plates with 10-15 embryos per well. For formate supplementation assays, 25 μM sodium formate (Sigma) was added. For nucleotide supplementation assays, 400 μM of AMP, UMP, GMP, IMP or TMP were added to embryos, or 200 μM of dA, dG, dU or T. 4-HNE (range of concentrations in ethanol) (Sigma) and Mtx (Sigma) (range of concentrations in DMSO) were added at 24hpf and refreshed every 24 hours.

### Aldehyde treatments

Stock solutions of fresh acetaldehyde and formaldehyde were made in a fume hood just before use. Various aldehyde concentrations were added to embryos kept in screw cap centrifuge tubes to limit aldehyde evaporation, and embryos scored for survival after 48 hours.

### RNA extraction and RT-qPCR

Samples to be processed for RT-qPCR were collected at the required stage and frozen on dry ice. RNA was extracted from frozen tissues with the Qiagen RNeasy Mini kit according to manufacturer’s instructions. RNA was quantified and quality checked using a Nanodrop 2000c (Thermo Scientific). 500 μg of RNA was used as input for Reverse Transcription using Superscript™ III reverse transcriptase (Invitrogen) and an Oligo(DT)_15_ primer (Promega). RT-qPCR was performed with Sybr Green® Lightcycler Green I Master mix (Roche), using a Lightcycler 480 instrument and associated software. *β-actin* was used as a housekeeping control (**Table S7**). Gene expression fold changes were found using the delta-delta ct method.

### Single cell sequencing experimental setup and sequencing

24hpf *Tg(mitfa:GFP; crestin:mCherry)* and were divided into groups of ∼500 embryos and treated with MoTP, and co-treated with either 10 μM CVT-10216 or equimolar DMSO. At 64hpf, MoTP was washed out, and embryos left to regenerate for 8 hours. Embryos were anaesthetized in MS-222 and trunks dissected, and a cell suspension of each treatment condition obtained as previously described (Manoli and Driever 2012). Samples were sorted on a FACSAria2 SORP instrument (BD Biosciences UK) as previously described (Brombin et al. 2021) but stage-matched non-fluorescent AB embryos also treated with MoTP used as a control to enable gating of *mCherry* and *GFP* fluorescence. 10,000 fluorescent cells were collected in 100 μl of 0.04% BSA/PBS. Single cell libraries were prepared using the Chromium Single Cell 3ʹ GEM, Library & Gel Bead Kit v3 (10x Genomics).

The samples were sequenced on a Nextseq 2000 using a P2 flow cell on a 100 cycle run. ∼2.97M reads passed quality filters for CVT-10216 treated, and ∼1.87M reads for DMSO-treated, however due to the greater number of cells processed in the CVT-10216 sample, the mean reads per cell were fairly equal (37,405-CVT vs 34,832-DMSO).

### Bioinformatics analyses

*Aldh2.2* expression within developmental melanocytes was visualized using the recent Brombin et al. GEO #: GSE178364 scRNA-seq dataset (Brombin et al. 2021).

For this study, FASTQ files were generated using CellRanger (v.3.1.0, 10x Genomics) mkfastq function with default settings and -qc parameter and aligned to the zebrafish STAR genome index using gene annotations from Ensembl GRCz11 release 94 with manually annotated entries for *GFP* and *mCherry*. Libraries were aggregated (CellRanger aggr pipeline) to generate a gene-barcode matrix. Gene matrices (13360 total, DMSO-5,394, CVT-7966), barcodes and features were uploaded to R (v. 4.0.5) and standard quality control filtering performed as previously described, to yield 4488 DMSO and 6795 CVT cells (Brombin et al. 2021). The dimensionality of the combined dataset was visualized with Elbow and JackStraw plots before running linear dimensional reduction. Louvain clustering was then performed using the FindNeighbors and FindClusters functions (dims=50, resolution=0.5) in Seurat (v. 4.0.3) (Hao et al. 2021). Data were projected onto 2D spaces using the same dimensions, using Uniform Manifold Approximation and Projection (UMAP). Cluster-specific genes were identified using Seurat as previously described (**Table S1,2**) (Brombin et al. 2021). Cluster calling was performed as previously described (Brombin et al. 2021) and by making unbiased pairwise comparisons based on gene overdispersion against published datasets GEO #: GSE131136 (Saunders et al. 2019) and NCBI SRA #: PRNJNA56410 (Farnsworth et al. 2020) and between the datasets presented in this paper as previously described (Brombin et al. 2021). Plots were generated either using Seurat or ggplot2 (v.3.3.5) (Wickham 2016). Prediction of cell cycle phase was performed with Seurat, using canonical cell cycle markers described in (Tirosh et al. 2016).

For DE analyses, scRNA-seq data were first corrected for zero-inflated counts by using the ZINB-WaVE package (v. 1.12.0) with default parameters (Risso et al. 2019). Then, the DEseq2 package (v. 1.30.1) (Love et al. 2014) was used to generate genelists of significantly (p.adj < 0.05) upregulated and downregulated genes (raw data in **Tables S3-6**). Pathway analyses were performed as previously described (Travnickova et al. 2019). GSEA analysis was performed using GSEA software (v. 4.1.0) with genelists generated through DeSeq2, using the “RunGSEAPreranked” function.

### Statistics

Statistical details of experiments and n numbers can be found in figure legends. Statistics and plots were generated using GraphPad Prism 7 (v. 7.0e) and R. Unless otherwise stated, experiments were replicated at least three times (N=3 biological replicates), with 10-15 embryos per condition.

## ACKNOWLEDGEMENTS

We are grateful to Cameron Wyatt and the IGC Zebrafish Facility for zebrafish management and husbandry, Elisabeth Freyer and the IGC FACS/10x facility, Ann Wheeler and the IGC Imaging Facility for supporting the imaging experiments, Richard Clarke at the Genetics Core ECRF for sequencing, and Jana Travnickova for sharing R code and expertise. JHP supported by US National Institutes of Health grants R01 OD011116 and R24 OD018555. EEP is funded by MRC HGU Program (MC_UU_00007/9), the European Research Council (ZF-MEL-CHEMBIO-648489), and Melanoma Research Alliance (687306).

## Author Contributions

Conceptualization: EEP, HB; Methodology: HB; Software: HB; Validation: HB; Formal analysis: HB, AB; Investigation: HB, AB, SP, JHP; Resources: SP, JHP, EEP; Writing original draft: EEP, HB; Writing review and editing: HB, AB, JHP, EEP; Visualization: HB, EEP; Supervision: JHP, EEP; Funding acquisition: JHP, EEP.

## SUPPLEMENTAL DATA

### Supplemental Figures

**Supplemental Figure S1:** *aldh2* expression in the McSC lineage, and generation of an *aldh2*−/− mutant line

**Supplemental Figure S2:** Results of aldehyde screen on *aldh2*−/− mutant zebrafish

**Supplemental Figure S3:** Identification of transcriptionally distinct scRNA-seq clusters during McSC regeneration

**Supplemental Figure S4:** Recovery of melanocyte regeneration following ALDH2i removal, and validation of Mtx treatment

### Supplemental Tables

**Supplemental Table S1:** scRNA-seq: top 30 cluster markers

**Supplemental Table S2:** scRNA-seq: metrics, clustering information and cell states

**Supplemental Table S3:** Differential expression analysis of *crestin+ mitfa-low* vs *crestin+ mitfa-high* cells

**Supplemental Table S4:** Differential expression analysis of *crestin+ mitfa-low* cells, DMSO vs CVT-10216 treated

**Supplemental Table S5:** Differential expression analysis of *crestin+ mitfa-high* cells, DMSO vs CVT-10216 treated

**Supplemental Table S6:** Differential expression analysis of iridophore cluster 9, DMSO vs CVT-10216 treated

**Supplemental Table S7:** Oligonucleotide sequences

### Supplemental Movies

**Supplemental Movie S1**: **McSCs generate progeny.**

Time-lapse video of a *Tg(crestin:mCherry;mitfa:GFP)* embryo during McSC regeneration (DMSO control). Embryos were treated with MoTP to kill differentiated melanocytes and initiate melanocyte regeneration. McSCs were followed for over 14 hours (post MoTP washout). McSCs are *crestin+ mitfa-low*, but then during or shortly after cell division, a cell strongly expresses GFP+ to become a *mitfa-high* cell, which then leaves the McSC compartment and migrates towards the epidermis.

**Supplemental Movie S2: ALDH2i inhibits McSCs ability to generate progeny.**

Time-lapse video of a *Tg(crestin:mCherry;mitfa:GFP)* embryo during McSC regeneration in the presence of ALDH2i. Embryos were treated with CVT-10216 to inhibit Aldh2, and co-treated with MoTP to kill differentiated melanocytes, and initiate melanocyte regeneration. McSCs were followed for over 14 hours (post MoTP washout, but in the presence of ALDH2i).

## SUPPLEMENTARY FIGURE LEGENDS

**Supplemental Figure S1:**
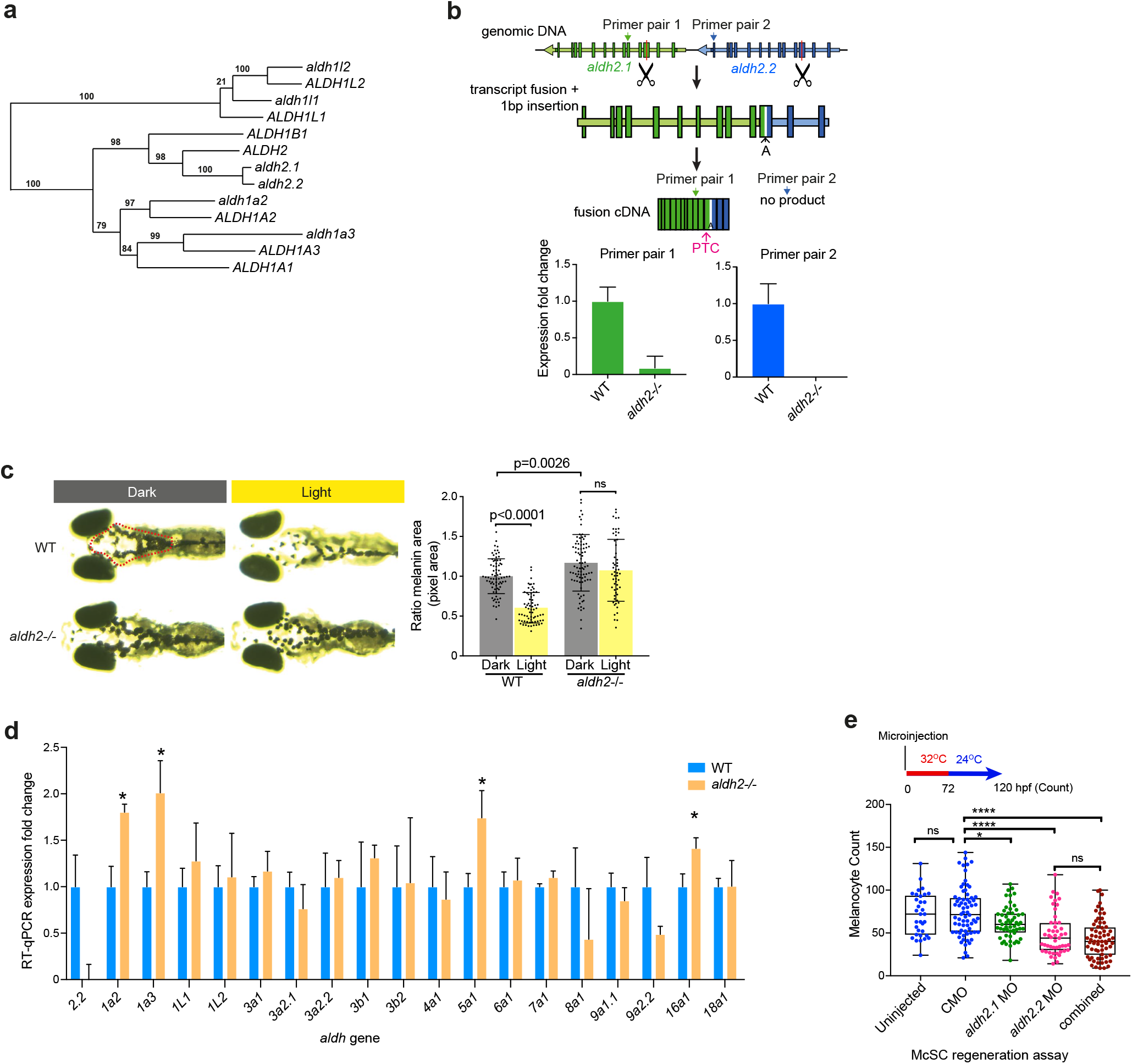
Aldh2 expression in the McSC lineage, and generation of an *aldh2−/−* mutant line. **a.** Phylogenetic tree showing the relationship between human *ALDH* and zebrafish *aldh* genes from the ALDH1 and ALDH2 families. **b.** sgRNAs targeting exon 3 of *aldh2.1* and *aldh2.2* were co-injected with Cas9. This caused deletion of the intergenic region and creation of a fusion transcript, with a base insertion and frame shift generating a premature stop codon (PTC). To confirm this deletion, RT-qPCR was performed with primers targeting Primer site 1, which should persist in the truncated fusion transcript, and Primer site 2 within the intergenic region. Relative expression of mutant fusion transcript in *aldh2−/−* mutants is shown relative to WT, and normalized to *β*-*actin*. N=3 biological replicates. Error bars represent SE. **c.** A camouflage response assay on WT or *aldh2−/−* embryos. Representative images are shown of embryos after adaptation to dark or light surroundings. Melanin coverage within the red outlined area was quantified. N=3 biological replicates. Error bars represent SD. P values indicated. Ordinary One-way ANOVA performed with Tukey’s multiple comparisons test. **d.** RT-qPCR showing quantification of wild type *aldh2.2* expression levels relative to other zebrafish *aldh* genes in 72hpf WT or *aldh2−/−* mutant embryos. *ß-actin* was used as a housekeeping control. N=3 biological replicates. Error bars show SE. Asterisks mark *aldh* genes upregulated >1.5-fold in *aldh2−/−* mutants compared to wild type. **e.** Regeneration assay on *mitfa^vc7^* embryos injected with 6 ng of standard control morpholino (CMO), or morpholinos against *aldh2.1, aldh2.2* or combination. Regenerated melanocytes are quantified. Each dot represents a single embryo. N=3 biological replicates. * p<0.0332, **** p<0.0001, ns: not significant. One-way ANOVA performed with Tukey’s multiple comparisons test.

**Supplemental Figure S2:**
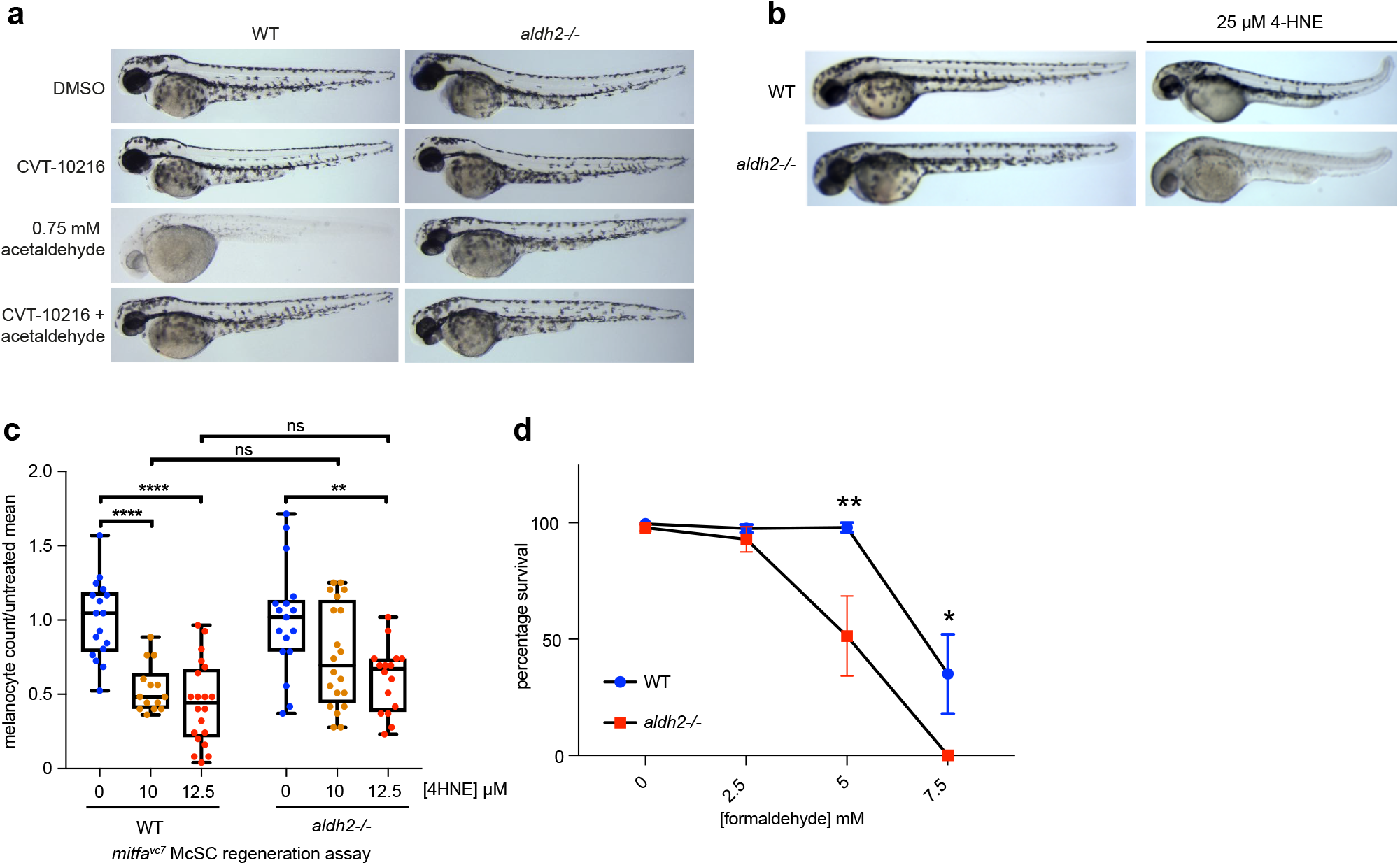
Results of aldehyde screen on *aldh2−/−* mutant zebrafish. **a.** Representative images of 72hpf WT and *aldh2−/−* mutant embryos treated with 0.75 mM acetaldehyde with or without CVT-10216. Unexpectedly, Aldh2 loss or deficiency confers resistance to acetaldehyde. N=3. **b.** Representative images of 72hpf WT and *aldh2−/−* mutant embryos treated with 25 µM 4-HNE, showing whole body sensitivity in the mutant. **c.** *mitfa^vc7^* melanocyte regeneration assay and subsequent quantification of embryos treated with increasing doses of 4-HNE, showing no significant difference between controls and *aldh2−/−; mitfa^vc7^* embryos in terms of reduction in regeneration potential after 4-HNE treatment. For comparison between genotypes, melanocyte numbers were normalized to the average untreated condition for each genotype. Each dot represents a single embryo. N=2. ** p<0.0021, **** p<0.0001, ns not significant. One way ANOVA with Tukey’s multiple comparisons. **d.** Survival percentage of WT and *aldh2−/−* mutant embryos treated with various concentrations of formaldehyde. N=6. Error bars are SE of the mean. ** p<0.0021. A two-way ANOVA with Sidak’s multiple comparisons test.

**Supplemental Figure S3:**
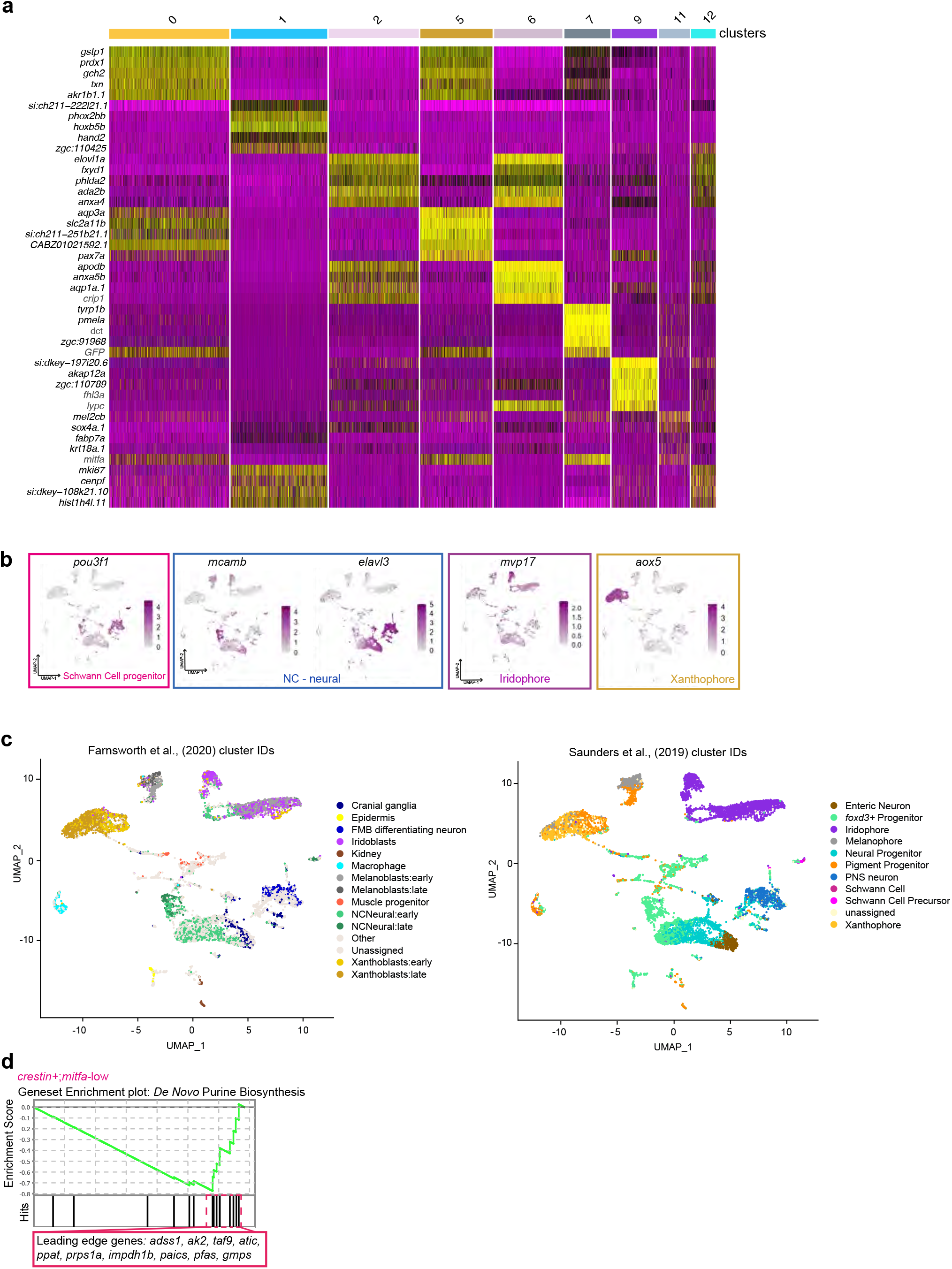
Identification of transcriptionally distinct scRNA-seq clusters during McSC regeneration. **a.** Heatmap showing top 5 cluster-defining genes per selected clusters. **b.** UMAP of the combined dataset showing gene expression of non-pigment clusters marked by *pou3f1* (Schwann Cell Progenitors), *mcamb* and *elavl3* marking NC-derived neural cells, *mvp17* marking iridophores, and *aox5* marking xanthophores. **c.** UMAP of this scRNA seq data mapped with cell identity annotation from Farnsworth et al (2020) and Saunders et al (2019). **d.** Enrichment plot of *de novo* purine biosynthesis signature upregulated in clusters 2,6,12 in CVT-10216 treated embryos compared to control, (NES -1.18, FDR <25%) Kolganov Smirnov test. Leading edge genes are listed.

**Supplemental Figure S4:**
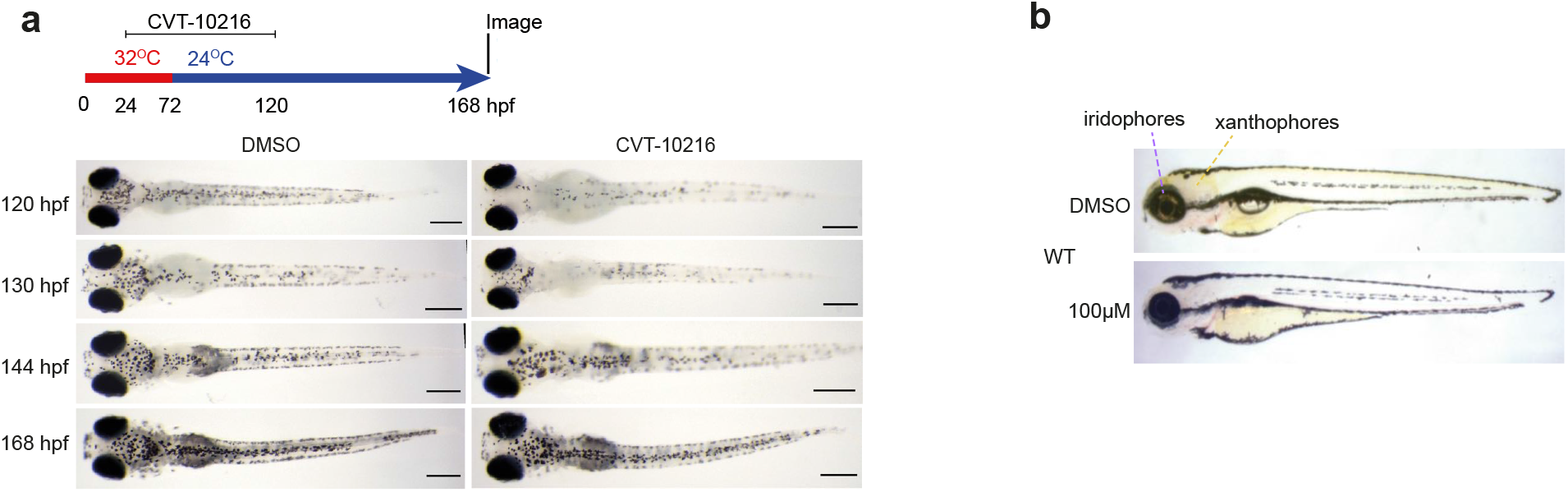
Recovery of melanocyte regeneration following ALDH2i removal, and validation of Mtx treatment. **a.** Extended regeneration assay on *mitfa^vc7^* embryos treated with CVT-10216 from 24-120hpf. After washout, larvae were imaged at a number of time points to monitor recovery/continuation of melanocyte regeneration. Representative images are shown from 5 embryos per condition. **b.** Mtx treated embryos (96 hpf) have lost reflective iridophore pigments (clearly observed in the eye) and yellow xanthophore pigments.

